# Variation in recombination rate affects detection of outliers in genome scans under neutrality

**DOI:** 10.1101/2020.02.06.937813

**Authors:** Tom R. Booker, Sam Yeaman, Michael C. Whitlock

**Author notes:** Author contributions: T.R.B., S.Y., and M.C.W. designed research; T.R.B. performed research; and T.R.B., S.Y., and M.C.W. wrote the paper.

## Abstract

Genome scans can potentially identify genetic loci involved in evolutionary processes such as local adaptation and gene flow. Here, we show that recombination rate variation across a neutrally evolving genome gives rise to mixed sampling distributions of mean *F*_*ST*_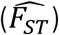, a common population genetic summary statistic. In particular, we show that in regions of low recombination the distribution of 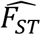 estimates have more variance and a longer tail than in more highly recombining regions. Determining outliers from the genome-wide distribution without taking local recombination rate into consideration may therefore increase the frequency of false positives in low recombination regions and be overly conservative in more highly recombining ones. We perform genome-scans on simulated and empirical *Drosophila melanogaster* datasets and, in both cases, find patterns consistent with this neutral model. Similar patterns are observed for other summary statistics used to capture variation in the coalescent process. Linked selection, particularly background selection, is often invoked to explain heterogeneity in 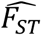 across the genome, but here we point out that even under neutrality, statistical artefacts can arise due to variation in recombination rate. Our results highlight a flaw in the design of genome scan studies and suggest that without estimates of local recombination rate, interpreting the genomic landscape of any summary statistic that captures variation in the coalescent process will be very difficult.

## Introduction

Genome scans have become a standard analysis in population genetics to identify particular regions that exhibit a significant departure from the background patterns observed in the rest of the genome. Many different statistics are used in such scans, and finding an outlier in the distribution of the test statistic of interest can indicate the presence of a causal variant affecting a phenotype, driving a selective sweep, or contributing to local adaptation, among other things (Cruickshank & Hahn, 2014; Haasl & Payseur, 2016; Hoban et al. 2016; Wolf & Ellegren, 2017). While some approaches are based on single-locus measures, a common practice is to aggregate statistics across loci within windows of the genome. As closely linked loci tend to share the same evolutionary history and therefore exhibit similar statistical signatures, this can potentially increase the power and decrease the influence of stochastic noise. However, because recombination rate can vary across the genome, some windows will tend to share evolutionary history among loci more strongly than other windows, and failure to account for this may result in bias in the estimation of the null distribution and testing procedure using window-based approaches. Here, we explore these issues using *F*_*ST*_ as an example, which is a statistic commonly used to search for signatures of local adaptation in the genome. There has been considerable discussion about how other processes, such as background or positive selection, can affect *F*_*ST*_, and a general acknowledgement of the importance of recombination variation for these alternative selection models (Burri, 2017; Cruickshank & Hahn, 2014; Nachman & Payseur, 2012), but little attention has been paid to the effects of recombination rate variation on the distribution of the neutral model. Given that all approaches that discuss selection and *F*_*ST*_ do so with reference to departures from a null model under drift, here we explore how recombination variation can also have critically important effects on the behaviour of the null model. This note emphasizes how failure to incorporate recombination rate into the null model leads to significant errors in our inferences and suggests more generally that this will also be important for any window-based approach to genome scans using other statistics.

Wright (1937) examined the behaviour of allele frequencies in metapopulations and defined *F*_*ST*_ as the correlation of allelic states between individuals from the same population. The evolutionary history of a population can be thought of as a series of genealogies describing the relationships between every individual at a certain point in the genome, and Slatkin (1991) showed that for a neutrally evolving metapopulation *F*_*ST*_ reflects the pattern of coalescence times of those genealogies. In particular, 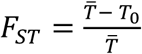, where *T*_0_ is the average coalescence time for a pair of alleles drawn from the same deme and 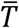 is the average coalescence time for a pair of alleles drawn from the metapopulation as a whole. *F*_*ST*_ varies across the genome; the mutations that happened to occur in the population and the individuals that happened to be sampled will give rise to variation in *F*_*ST*_ at different points in the genome. There are numerous methods to average *F*_*ST*_ across linked sites in an attempt to account for these sources of variation (Bhatia et al. 2013; Holsinger & Weir, 2009). Furthermore, the underlying demographic process is also stochastic and will generate different histories across the genome giving rise to heterogeneity in *F*_*ST*_.

In the context of genome scans, 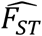 is often calculated in analysis windows of a fixed physical size or number of single nucleotide polymorphisms (SNPs), and the genome-wide distribution is examined under the implicit assumption that all windows share the same sampling distribution. This assumption is reflected by the discrete thresholds that are often applied to genome-wide distributions of 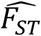 to identify outliers in empirical studies (Harpak et al., 2020; Lai et al., 2019; Ma et al., 2017; Olofsson et al., 2019; Talla et al., 2019; Zhou et al., 2019). However, recombination breaks down associations across genetically linked sites allowing coalescence times to vary across the genome, causing some windows to reflect more or fewer coalescent histories than others. Thus, the sampling distribution of 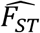, and indeed any summary statistic that captures aspects of coalescent history, will likely covary with the rate of recombination.

Consider a 10,000bp analysis window in a neutrally evolving genomic region that experiences virtually no recombination (for example, the mitochondrial genome or non-pseudoautosomal regions of the Y-chromosome in humans). Every nucleotide within the window would share a single genealogy, and every polymorphism present would reflect exactly the same coalescent history. If there were a large number of polymorphisms in the window, the combined 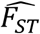 across all of them may provide a very precise estimate of their shared history. If one had a large number of these analysis windows and examined the distribution of their averages, it should resemble the coalescent distribution of *F*_*ST*_, which, for a population conforming to the island model, can be approximated using the *χ*^2^ distribution (Lewontin & Krakauer, 1973), which has a long upper tail.

Now consider the alternate extreme, 10,000bp of freely recombining nucleotides. In this case, all sites would have a distinct evolutionary history, each representing a quasi-independent instantiation of the coalescent process. As a result, every polymorphism present would carry information on a different genealogy, and each of these may have a distinct *F*_*ST*_. The observed 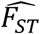 across all polymorphisms in this set provides an estimate of the mean of the coalescent distribution of *F*_*ST*_. If one had a large number of these sets of loci, the values of 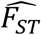 should be tightly distributed about the mean, and central limit theorem predicts that the distribution should be approximately Gaussian.

Obviously, 10,000 contiguous nucleotides do not experience free recombination, but eukaryotes do exhibit wide variation in recombination rates across their genomes. For example, humans (Kong et al. 2010), collared flycatchers (Kawakami et al. 2014) and *Drosophila melanogaster* (Comeron et al. 2012) all exhibit recombination rate variation over at least an order of magnitude across substantial proportions of their genomes (Supplementary Figure 1). If the sampling distribution of 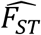 had a longer tail in regions of the genome with low recombination and was narrower in more highly recombining ones, heterogeneous landscapes may arise as a statistical artefact under neutrality. Determining outliers from the genome-wide distribution of summary statistics may, therefore, result in false positives and negatives depending on the local recombination rate. In this study, we demonstrate that the sampling distribution of 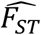 does indeed depend on the local recombination rate and show how this may affect genome-scans using simulated and empirical datasets.

## Results and Discussion

To determine whether the sampling distribution of 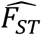 is affected when recombination rates vary over orders of magnitude, we simulated genomic data under an island model of population structure. Different distributions of mean 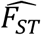 arose under different recombination rates (Figure 1). The theoretical expectation of *F*_*ST*_ was the same in all of the simulations shown in Figure 1, and the average 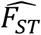 was very close to the expectation in all cases. However, low recombination rates led to longer-tailed distributions of 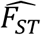 and higher rates led to distributions that were tighter about the mean (Figure 1). Similar patterns are seen under a variety of demographic models (Supplementary Figure 2).

**Figure 1.**
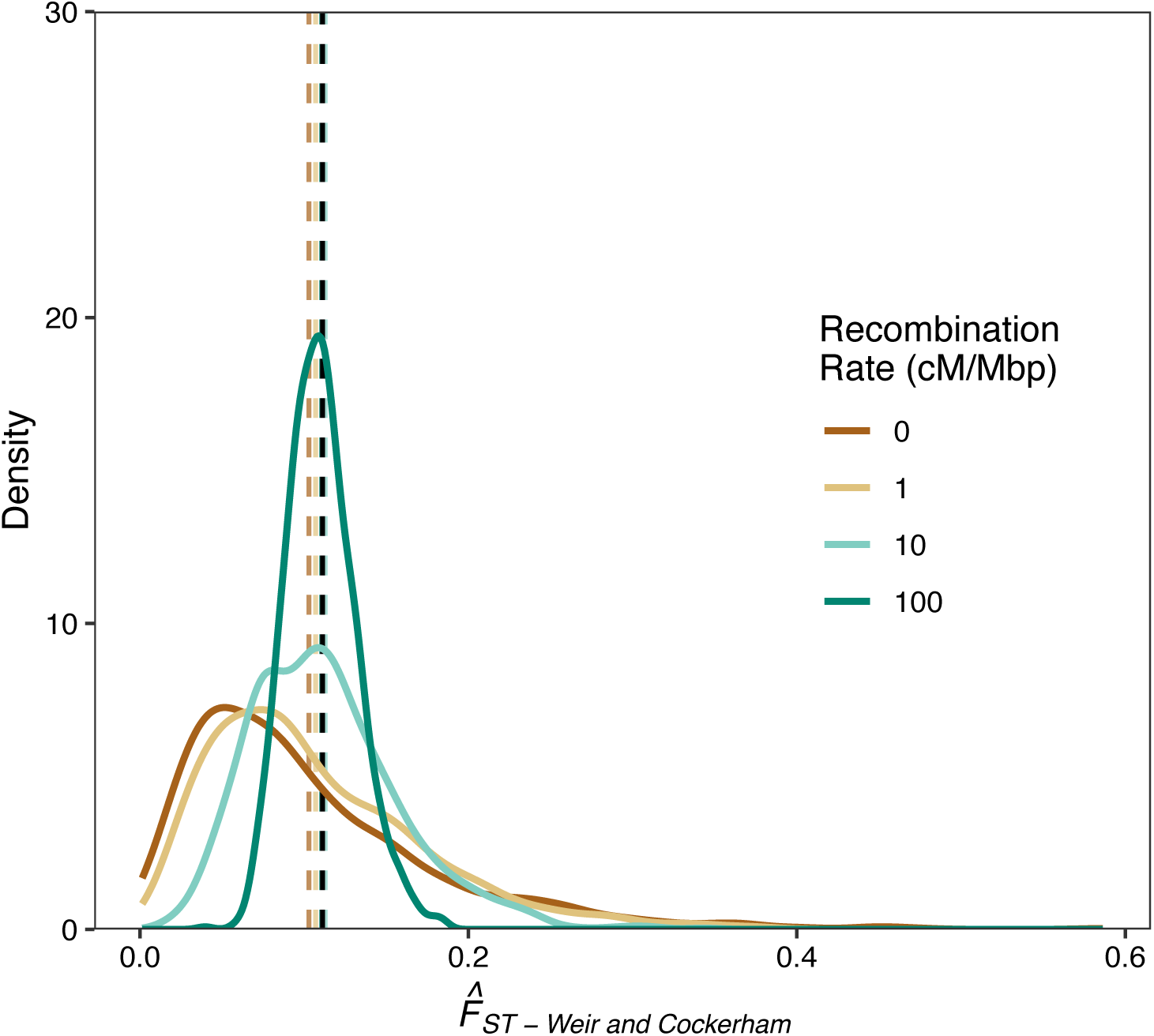
The distribution of *F*_*ST*_ calculated in 10 kbp windows for an island population with *N*_*e*_ = 100 haploids per deme. *F*_*ST*_ was averaged using the method of Weir and Cockerham (1984). The dashed vertical line is the theoretical expectation of *F*_*ST*_ for the simulated population, and the coloured vertical lines indicate the means for each set of simulations.

A typical method for determining outliers in genome scans, used by all the empirical studies cited in the introduction, is to examine the top *n*^*th*^ percentile of a particular summary statistic. If one were performing a genome scan in a species that exhibited recombination rate variation even over a single order of magnitude, one would expect the tail of the genome-wide distribution of 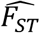 to be enriched for analysis windows in low recombination regions under neutrality (Figure 1). Indeed, the recombination rates shown in Figure 1 obviously lead to different distributions of 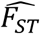, but the extent to which this will affect the analysis of real organisms will depend on the pattern of variation in recombination rates across their genomes.

We examined the relationship between 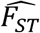 and recombination rate across the *D. melanogaster* genome using simulated and empirical datasets. We simulated the entire *D. melanogaster* genome under a model of isolation-with-migration incorporating recombination rate variation using *stdpopsim* (Adrion et al. 2019) and performed genome scans on the resulting data (Figure 2). We performed a similar genome scan on the population genomic data from North American *D. melanogaster* populations generated by Reinhardt et al. (2014). In both cases, we found a significant enrichment of outliers at low recombination rates compared to high recombination rates when we defined outliers based on the 95^th^ percentile of 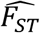 (simulated data Fisher’s exact test *p*-value < 10^−15^; empirical data Fisher’s exact test *p-*value = 0.00265). Furthermore, both the simulated and empirical datasets exhibited significant negative correlations between the variance of 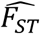 and recombination rate (Figure 2C; simulated data, Kendall’s *τ* = −0.746, *p*-value < 10^−15^; empirical data, Kendall’s *τ* = −0.193, *p*-value = 0.00445). We found qualitatively similar patterns in our simulated data when analysis windows were based on a fixed number of SNPs (Supplementary Figure 3). Of course, the genomic landscape of 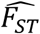 in North American *D. melanogaster* has likely been shaped by processes other than just migration and genetic drift. Indeed, Figure 2A shows at least one 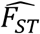 outlier at high recombination rates, which our simulations suggest would be unlikely under neutrality (Figure 2B). In summary, our findings suggest that applying fixed thresholds to the genome-wide distribution of 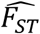 may enrich for false positives in low recombination regions and be overly conservative in highly recombining regions of the genome.

**Figure 2.**
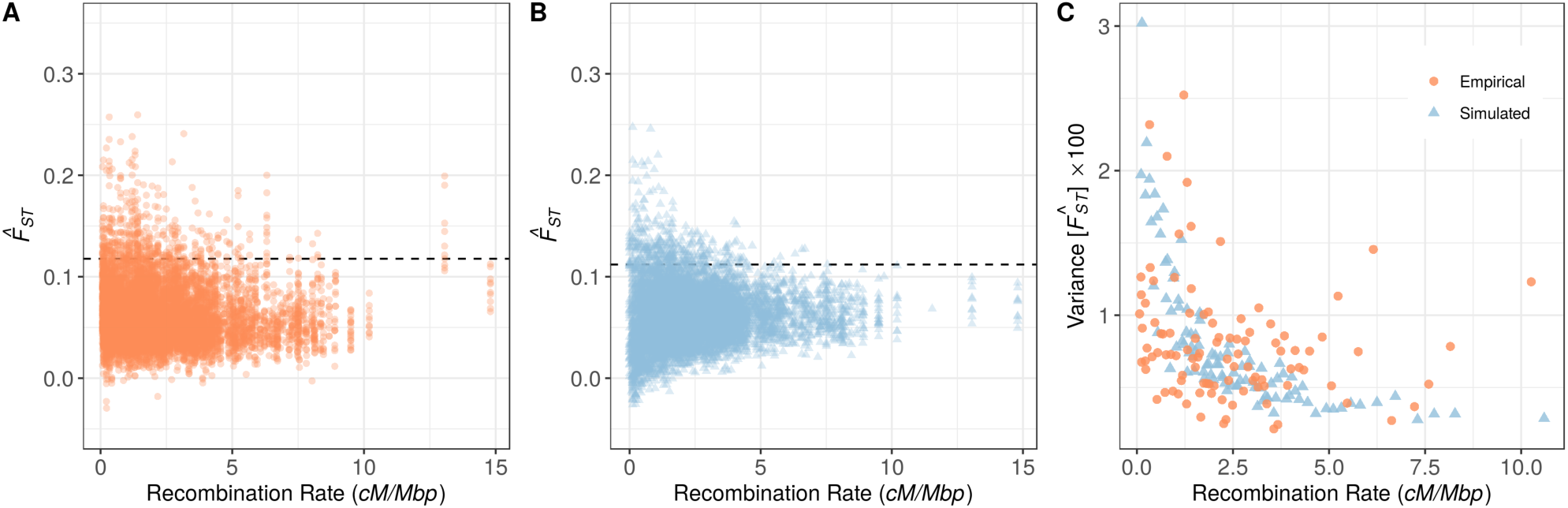
The plot of 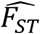 calculated in 10,000bp analysis windows against local recombination rate for *Drosophila melanogaster* from A) the North American populations of *Drosophila melanogaster* analysed by Reinhardt et al. (2014) and B) coalescent simulations of the genome under a model of isolation with migration. C) Shows the variance in 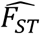 against recombination rate for both empirical and simulated datasets. The horizontal lines in A and B are the 95th percentiles of the genome-wide distribution in each case

Previous studies have shown greater heterogeneity in *F*_*ST*_ in regions of low recombination rate across the genome, but these have interpreted their findings purely in terms of linked selection (e.g. Duranton et al. 2018; Gagnaire et al. 2018; Samuk et al. 2017). Indeed, the effects of selection at linked sites, particularly background selection, have been invoked to explain the role of recombination in generating patterns of variation in *F*_*ST*_, and other summary statistics across the genome, in particular the so-called “islands of differentiation” involved in speciation (Adrion et al., 2019; Burri, 2017; Cruickshank & Hahn, 2014; Nosil et al., 2009). In this study, we are suggesting that genome-scans using analysis windows should incorporate recombination rate variation into the null model used to interpret the data, even in the absence of selection. Obviously, linked selection, which has the greatest impact when recombination rates are low, should be considered as well, but without considering the statistical artefacts that arise under neutrality, it will be difficult to distinguish pattern and process from genome-scan data.

It has long been known that the variance of summary statistics such as the number of segregating sites responds to variation in recombination rate (Wakeley, 2009), so the difficulty of comparing analysis windows of discrete physical size across varying recombination rates will extend to other data summaries that capture variation in the coalescent process. For example, genome-scan studies may analyse nucleotide diversity between populations (*D*_*XY*_), which is proportional to *T*_*B*_, the coalescence times between populations, within population diversity (*π*_*w*_), Tajima’s *D* (Tajima, 1989) or the haplotype based statistics developed by Garud et al. (2015), for instance H12. We calculated each of those statistics from our simulated data and, as expected, the variance of each increased with recombination rate (Supplementary Figure 4). Genome-scan studies often examine combinations of summary statistics that respond differently to various evolutionary processes, but even under neutrality extreme values for multiple summary statistics may occur (Supplementary Figure 5). Without an understanding of what constitutes an extreme value of a given summary statistic for a particular recombination rate, interpreting the genome-wide distribution may be difficult.

There are obviously cases, however, where the heterogeneity in genome-scans introduced by recombination rate variation would not affect the ability to determine the loci involved in evolutionary processes. When there is a monogenic or oligogenic architecture of adaptation or speciation, the regions involved may clearly stand out from the genomic landscape of summary statistics. For instance, Jones et al. (2018) performed a genome-scan comparing snowshoe hares with white and brown winter coats and found an analysis window containing the *Agouti* gene had by far the highest between-population divergence genome-wide. However, in their study, Jones et al. (2018) also drew on orthogonal evidence from a genome-wide association study and gene expression data to support their conclusions about that region’s involvement in adaptation. In cases where processes such as local adaptation involve a highly polygenic architecture, the effects on genome-wide variation may be more subtle and difficult to interpret (Latta, 1998; Le Corre & Kremer, 2012; Yeaman, 2015).

The results from this study should emphasize the need to incorporate recombination into the null model used to interpret genome scans. Generating recombination maps is a large undertaking, and there are numerous systems in which it is not feasible to obtain them, but new methods for estimating recombination rate may greatly facilitate this (Dreau et al. 2019; Sun et al. 2019). In cases where accurate recombination rate estimates are not available, once could perhaps determine analysis windows based on patterns of average linkage disequilibrium across sites. Alternatively, if accurate recombination rate estimates were available, outlier thresholds could be determined for discrete recombination rate bins, or analysis windows could be designed using fixed genetic, rather than physical, distances. Further work is required to develop and test analysis methods that are robust to recombination rate variation. Without accounting for recombination rate variation in some way, however, it will be difficult to interpret variation in the genomic landscape of summary statistics.

## Materials and Methods

Coalescent simulations of structured populations were performed using *msprime* v0.7.3 (Kelleher et al. 2016). We simulated an island model of 100 demes, each with *N*_*e*_ = 100 haploid individuals. Migration was constant between all demes and was set to *M* = *N*_*e*_*m* = 2, giving an expected *F*_*ST*_ of 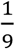. After simulating a particular population, we added mutations to the resulting tree sequence at a rate of 10^−6^. We extracted 100 individuals each from two demes then calculated the following summary statistics using *scikit-allel* v1.2.1: the weighted average of *F*_*ST*_ using the method of Weir and Cockerham (1984); between population genetic diversity (*D*_*XY*_); Tajima’s *D* (Tajima, 1989); and Garud’s H12 (Garud et al. 2015). For Garud’s H12, we only examined the first 400 SNPs present in the simulated regions. We added mutations and calculated summary statistics 100 times for each simulated population and took the average across replicates. We performed 1,000 such simulations for each recombination rate tested. Smoothed density plots were generated with default settings in *ggplot2* (Wickham, 2016).

We re-analysed allele frequency data from the North American *Drosophila* populations generated by Reinhardt et al. (2014). In that study, pooled sequencing was used to estimate derived allele frequencies for pools of 16 isofemale lines each from Maine and Florida. To obtain accurate allele frequencies we excluded any SNP that was reported at a depth greater than or less than one standard deviation from the mean coverage.

We simulated the entire *Drosophila melanogaster* genome under a model of isolation with migration in *stdpopsim* v0.1.1. An ancestral population with effective size 2*N*_*e*_ = 344,120 individuals, split into two demes of *N*_*e*_ individuals *N*_*e*_ generations in the past. Symmetrical migration of 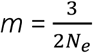 migrants per generation occurred after the split. The migration rate was chosen to approximately match simulated *F*_*ST*_ with the empirical data. We sampled 20 simulated haploid individuals from each deme.

We analysed the genome-wide distribution of *F*_*ST*_ in the empirical and simulated *Drosophila melanogaster* datasets. Because the Reinhardt et al. (2014) data was generated using pooled-sequencing, we calculated *F*_*ST*_ for each polymorphism and averaged across sites using the formulae for haploids given in Weir (1990). 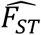 was calculated in 10,000 bp non-overlapping windows using the ratio-of-averages approach. Recombination rates from the Comeron et al. (2012) map were obtained using the *Drosophila* recombination rate calculator (http://petrov.stanford.edu/cgi-bin/recombination-rates_updateR5.pl; Fiston-Lavier et al. 2010). For both the simulated and empirical *D. melanogaster* data, we only examined the “normally recombining” regions of the *Drosophila* genome, following Reinhardt et al. (2014). We divided the genome into 75 equally sized bins based on recombination rate. We calculated Kendall’s correlation between the variance of 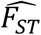 and the mean recombination rate in each bin. We classified analysis windows as outliers if 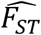 was greater than the 95^th^ percentile genome-wide. We examined the analysis windows with the 1,000 largest and smallest recombination rate estimates and used Fisher’s exact test to determine whether there was a significant enrichment of outliers in low recombination regions. We excluded analysis windows from the simulated data that had recombination rates of 0.

## Acknowledgements

Thanks to Sally Otto, Andrea Thomaz and Wouter van der Bijl for helpful discussions and to Reto Burri, Katie Lotterhos and Matt Pennell for comments on the manuscript. Thanks to Andrew Kern for insights and for providing the *Drosophila* data and to Takeshi Kawakami for providing the flycatcher recombination rate estimates. This study is part of the CoAdapTree project which is funded by Genome Canada (241REF), Genome BC and 16 other sponsors (http://coadaptree.forestry.ubc.ca/sponsors/). MCW is supported by a Discovery Award from NSERC. SY is supported by a Discovery Award from NSERC and a research chair from Alberta Innovates.

## Supplementary Material

**Supplementary Figure 1.**
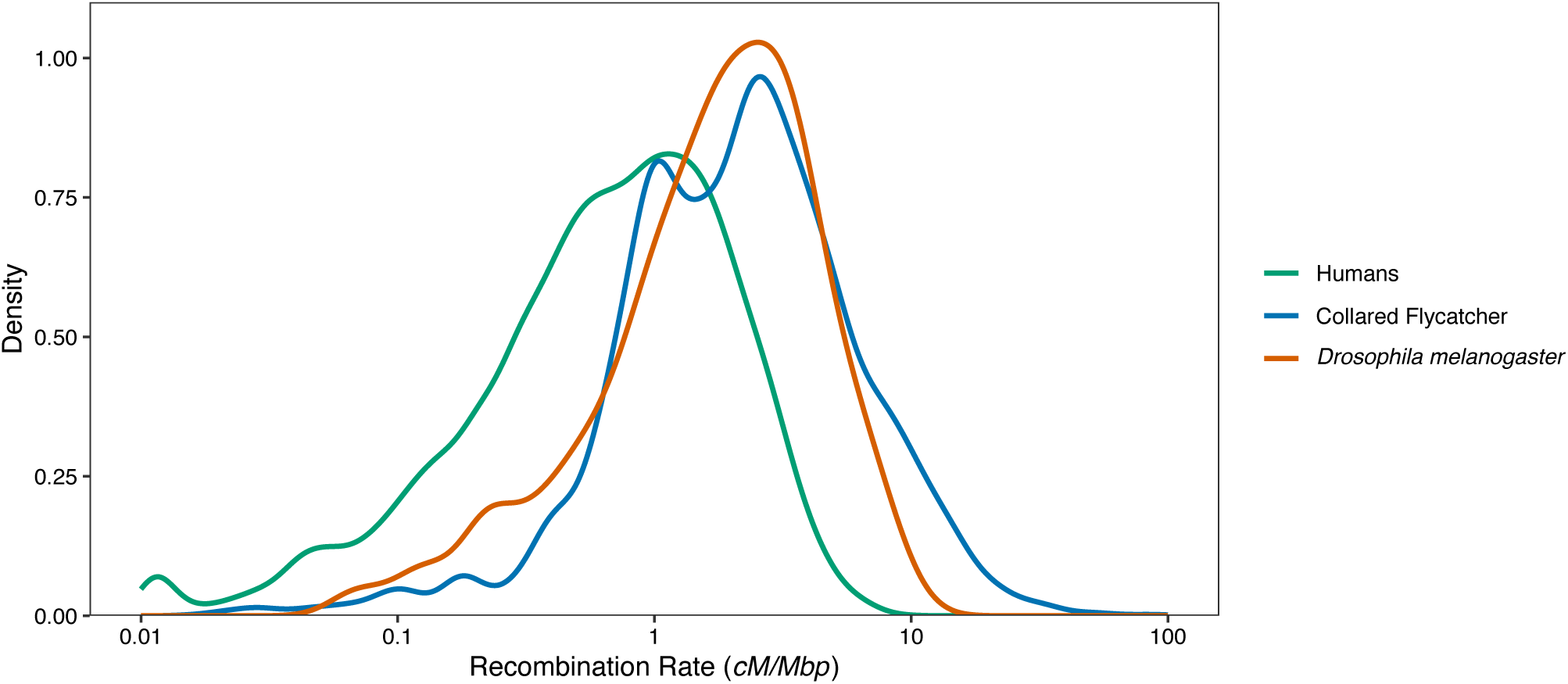
Recombination rate variation in three animal species calculated for 200 kbp genomic windows. The human map comes from Kong et al. (2010)(data downloaded from https://www.decode.com/addendum/). The collared flycatcher data come from Kawakami et al. (2014), and the *Drosophila melanogaster* data come from Comeron et al. (2012) and were obtained using the *Drosophila* recombination rate calculator (http://petrov.stanford.edu/cgi-bin/recombination-rates_updateR5.pl; Fiston-Lavier et al. 2010)

**Supplementary Figure 2.**
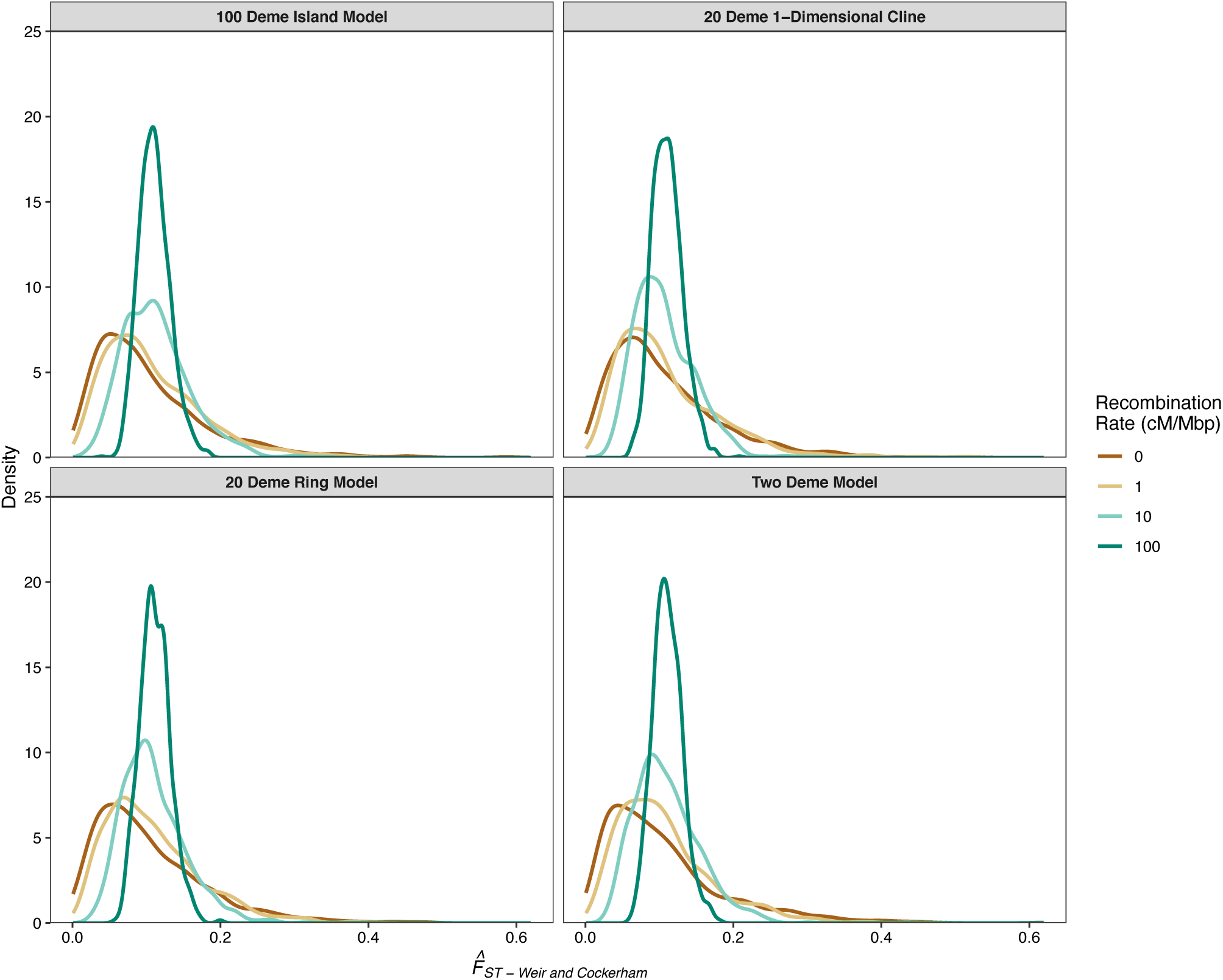
The distribution of *F*_*ST*_ calculated in 10 kbp windows for four differently structured metapopulations. *F*_*ST*_ was averaged using the method of Weir and Cockerham. Note, the 100-deme island model is represented in Figure 1 in the main text.

**Supplementary Figure 3.**
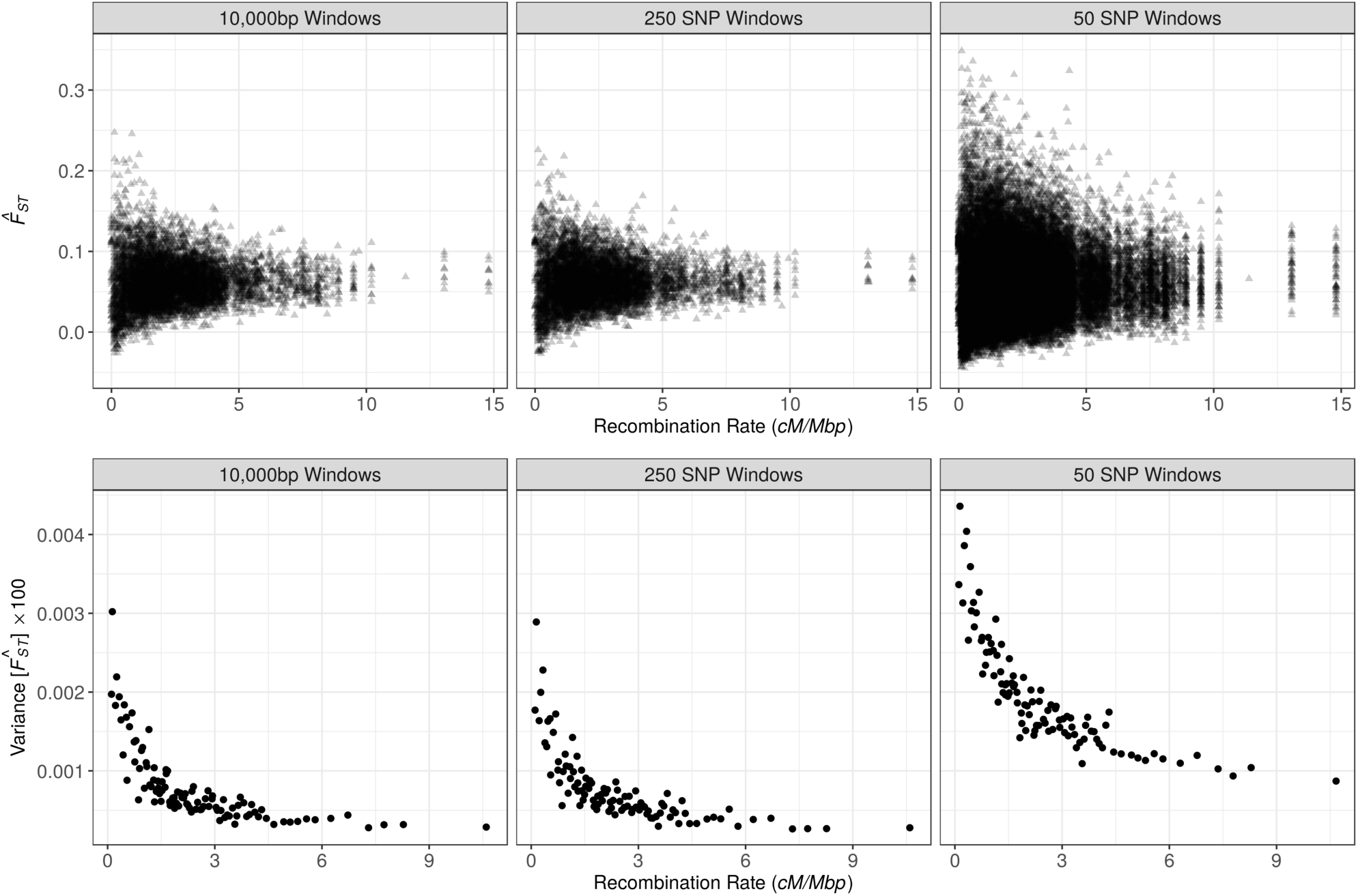
The plot of 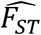 and the variance in 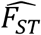 against local recombination rate for coalescent simulations of the genome under a model of isolation with migration. The panels show 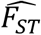 calculated in analysis windows of either a fixed physical size (10,000 bp) or fixed number of SNPs (250 or 50).

**Supplementary Figure 4.**
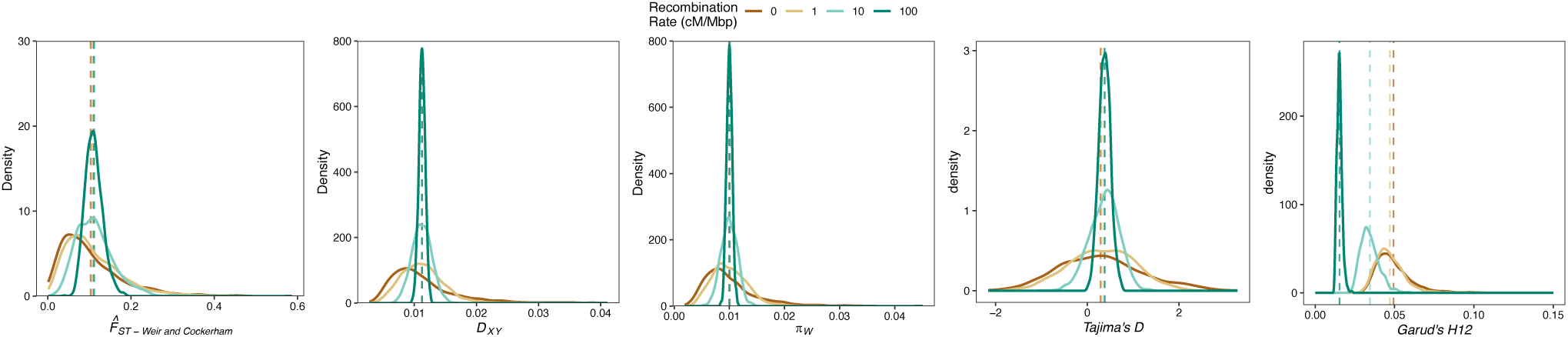
The distribution of five population genetic summary statistics often used in genome scan studies. All statistics were calculated in 10 kbp windows except for H12 which was calculated using a window of 400 SNPs. *F*_*ST*_ was averaged using the method of Weir and Cockerham. Populations were simulated according to a 100-deme island model. Vertical lines indicate the mean of a given distribution.

**Supplementary Figure 5.**
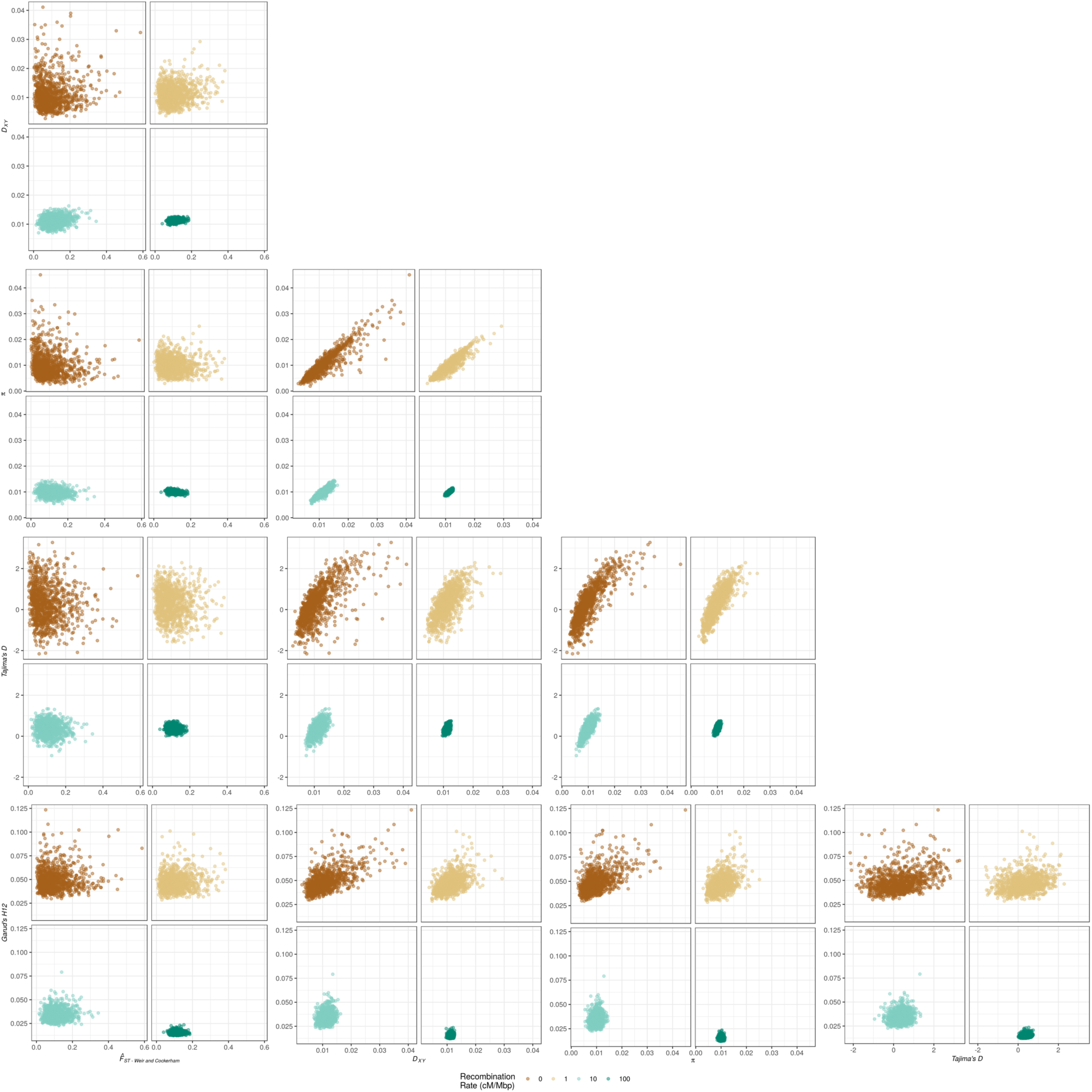
Pairwise comparison of five population genetic summary statistics often used in genome scan studies under of four recombination rates. Each panel shoes the results from 1,000 independent coalescent simulations. All statistics were calculated in 10 kbp windows except for H12 which was calculated using a window of 400 SNPs. *F*_*ST*_ was averaged using the method of Weir & Cockerham, (1984). *π* and *D*_*XY*_ are expressed per base pair. Populations were simulated according to a 100-deme island model.

